# Genetic diversity of pangolin coronaviruses reveals a key immuno-evasive substitution at spike residue 519

**DOI:** 10.1101/2025.04.01.646621

**Authors:** Maximilian Stanley Yo, Yu Kaku, Yusuke Kosugi, Jarel Elgin Tolentino, Yunlong Cao, Kei Sato

## Abstract

Malayan pangolins are unprecedented hosts for several SARS-CoV-2-related coronaviruses, which have previously been known to only infect *Rhinolophus* bats. Much debate has hence surrounded their possible role as intermediate hosts in the emergence of SARS-CoV-2, but the virological phenotypes of most pangolin coronaviruses (pCoVs) remain unclear. Here, we comprehensively analyze all pCoVs to date identified from trafficked pangolins seized in the Guangdong province of China, which are remarkably similar to SARS-CoV-2 in the spike protein. We explore an unknown genetic diversity within these viruses and uncover how this diversity translates to different virological phenotypes. Strikingly, several Guangdong pCoVs harbor a lysine substitution at residue 519 of spike protein, which contributes to marked immune evasion potentially by modulating the conformational state of spike protein. Furthermore, we highlight that a similar immuno-evasive mutation at residue 519 of the spike protein was acquired by SARS-CoV-2. These findings support that pangolin– and human-infecting coronaviruses represent independent spillover events from natural bat reservoirs, and that immuno-evasive mutations at residue 519 may be a common vector of viral evolution in coronaviruses that infect non-bat hosts.

## Introduction

Severe acute respiratory syndrome coronavirus 2 (SARS-CoV-2) emerged at the end of 2019 and caused the coronavirus disease 2019 (COVID-19) pandemic. This global event drew much attention to coronavirus surveillance (1), which has yielded many viruses related to SARS-CoV-2. Independently to SARS-CoV-2, SARS-CoV also caused outbreaks mainly in East and Southeast Asian countries in 2002 (2). Both SARS-CoV and SARS-CoV-2 are classified into the family *Coronaviridae*, the genus *Betacoronavirus*, and the subgenus *Sarbecovirus* (3). Sarbecoviruses have been found to primarily infect bats belonging to the genus *Rhinolophus* (4–6), building on a long-standing consensus that *Rhinolophus* bats are a natural reservoir of sarbecoviruses (7). However, in recent years a handful of sarbecoviruses have also been detected in smuggled Malayan pangolins (*Manis javanica*) in Southern China, specifically in the Guangdong (8–12) and Guangxi (11,12) provinces.

Malayan pangolins are distributed across much of Southeast Asia, sharing a common ecological niche and distribution with *Rhinolophus* bats (13). The proximity of Malayan pangolins to *Rhinolophus* bats in the wild places them in an environment where spillover of coronaviruses could occur. For instance, sarbecoviruses have been detected in *Rhinolophus* bats in Cambodia, geographically overlapping with pangolin populations (14). Furthermore, following the identification of pangolin coronaviruses (pCoVs) in Southern China, SARS-CoV-2 neutralizing antibodies were identified in a pangolin in Southern Thailand (15) and in a pangolin carcass likely originating from Indonesia (16), suggesting that spillover events of coronaviruses into pangolin populations may occur naturally in Southeast Asia.

Malayan pangolins are among the most trafficked species globally (13). The illegal wildlife trade has allowed pangolins to act as a “vehicle” for various viruses (including coronaviruses) to cross country borders (17). As pangolins encounter wildlife smugglers, there is a risk that pCoVs may spill into human populations, causing future coronavirus disease emergence. Indeed, all pCoVs to date were identified using samples from seized pangolins of unknown country of origin in anti-smuggling operations (9–12).

The Guangdong pCoVs (GD pCoVs) were identified using samples from up to nine pangolins in March 2019 (9–11) and one pangolin in July 2019 (10), all sampled at the Guangdong Wildlife Rescue Center, China. On the other hand, the Guangxi pCoVs (GX pCoVs) were identified using samples originating from 5 pangolins seized by Guangxi Customs from August 2017 to January 2018 (12). The sequenced pCoVs were found to be highly similar to SARS-CoV-2, leading to debate surrounding whether pCoVs carried by smuggled pangolins played a role in the emergence of SARS-CoV-2 (18,19).

Previous studies into the pCoVs found that the virological phenotypes of pCoVs resemble those of SARS-CoV-2. For instance, similarly to SARS-CoV and SARS-CoV-2, both GD and GX pCoVs can use both pangolin and human angiotensin converting enzyme 2 (ACE2) as a functional receptor for infection (20–22). Additionally, certain GD pCoV strains are susceptible to neutralization by human anti-SARS-CoV-2 humoral immunity induced by SARS-CoV-2 infection and vaccination (22,23). However, although several GD pCoV sequences identified from different pangolin individuals have been published (9–11), the virological phenotypes of most pCoVs remain unknown. In this study, we characterize the genotypic and phenotypic diversity of all GD pCoVs and analyze how these characteristics inform the evolution of sarbecoviruses in Malayan pangolins.

## Results

### The GD pCoV RBD is closely related to the SARS-CoV-2 RBD

We first described the phylogenetic relationship of pCoVs to other sarbecoviruses. A number of sarbecoviruses have been detected in *Rhinolophus* bats, but as of April 2025, only 16 pCoV whole genome or spike (S) sequences, including 11 GD pCoVs and 5 GX pCoVs, have been published (9–11). We constructed three phylogenetic trees, which are based on the sequences of the whole genome (**Fig. 1A**), S protein (**Fig. 1B**), and S receptor-binding domain (RBD) (**Fig. 1C**). The strains and accession numbers are summarized in **Table S1**. All trees showed that all pCoVs are phylogenetically closer to SARS-CoV-2 (strain Wuhan-Hu-1) than SARS-CoV (strain Tor2) (**Fig. 1A–C**). Consistent with previous reports (12,24), pCoVs are phylogenetically separated into two clades: GD pCoVs and GX pCoVs (**Fig. 1A–C**). Moreover, the molecular phylogenetic tree of the S RBD showed that all eleven GD pCoVs are more closely related to SARS-CoV-2 than GX pCoVs (**Fig. 1C**).

**Fig. 1.**
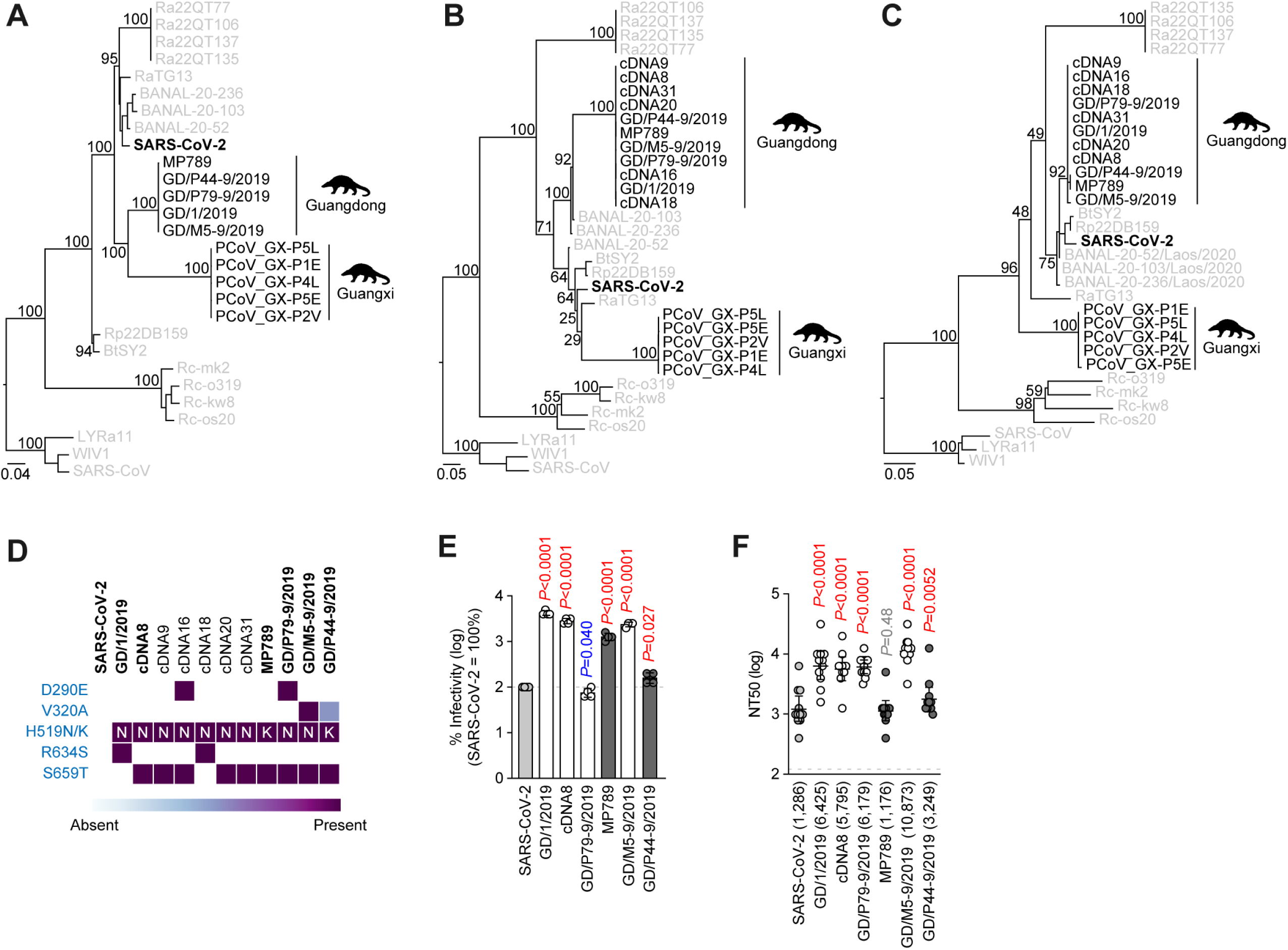
Phylogenetic relationship of GD pCoVs and differing sensitivity to human humoral immunity induced by SARS-CoV-2 vaccine. (**A–C**) Maximum likelihood trees of SARS-CoV-2 related pangolin coronaviruses, SARS-CoV-2 (strain Wuhan-Hu-1), and other sarbecoviruses. The trees based on the whole genome sequences (**A**), S nucleotide sequences (**B**), and S RBD amino acid sequences (**C**) are shown. SARS-CoV-1 (strain Tor2) and two SARS-CoV-related coronaviruses (strains WIV1 and LYRa11) are included as an outgroup. SARS-CoV-2 is shown in bold. GD pCoVs and GX pCoVs are shown in black, and annotated based on previous published work (9–12). Other sarbecoviruses are shown in gray. Node support is shown above key branches. Scale bar indicates genetic distance (nucleotide/amino acid substitutions per site). (**D**) Amino acid substitutions in the S proteins of all eleven GD pCoVs detected to date. Substitutions and site positions are described in reference to SARS-CoV-2. The sequence of GD/P44-9/2019 has an ambiguous nucleotide resulting in a possible translation to alanine or valine at position 320. Since the alanine translation makes GD/P44-9/2019 identical to MP789, the valine translation at this position was adopted. The viruses in bold were used in this study. The substitutions in GD pCoVs and their surrounding amino acids are summarized in **Fig. S1** (**E**) Pseudovirus assay of GD pCoVs. HIV-1-based reporter viruses pseudotyped with the S proteins of GD pCoVs were prepared. The pseudoviruses were inoculated into HOS-TMPRSS2 cells stably expressing human ACE2. The mean and SD of the log-transformed percent infectivity compared to SARS-CoV-2 are shown (representative of biological triplicate). The horizontal dashed line indicates the infectivity of SARS-CoV-2. Assays were performed in quadruplicate. Statistically significant differences versus SARS-CoV-2 were determined by the two-sided Student’s t test and *P* values are shown in the figure. (**F**) Neutralization assay of GD pCoVs. Neutralization assays were performed with pseudoviruses harboring the S proteins of GD pCoVs and SARS-CoV-2 against sera elicited by 3-dose Pfizer-BioNTech COVID-19 vaccination (n = 11). Assays for each serum sample were performed in triplicate to determine the 50% neutralization titer (NT50). Each point represents one NT50 value, and the geometric mean and 95% confidence intervals are shown. The number in parentheses indicates the geometric mean of NT50 values. The horizontal dashed line indicates the limit of detection (120-fold). Statistically significant differences versus SARS-CoV-2 were determined by the two-sided Student’s t-test paired by serum sample and *P* values are shown in the figure. Information on the vaccine sera donors is summarized in **Table S3**. In (**E**) and (**F**), open circles/columns represent viruses that harbor an asparagine at position 519, while filled circles/columns represent viruses that harbor a lysine at position 519. SARS-CoV-2 is included as a control and is shaded in gray. Red and blue values indicate a significant increase and decrease compared to SARS-CoV-2, respectively.

### GD pCoVs exhibit phenotypic diversity in infectivity and susceptibility to humoral immunity induced by SARS-CoV-2 vaccination

To compare the virological phenotypes of different GD pCoV strains, we compared all eleven of the publicly available GD pCoV S sequences. The sequences consist of three whole genomes [BetaCoV/pangolin/Guangdong/1/2019 (GD/1/2019) (9), MP789 (10), and GD/P79-9/2019 (11)], two partial genomes (GD/M5-9/2019 and GD/P44-9/2019) (11), and six S sequences (cDNA8, cDNA9, cDNA16, cDNA18, cDNA20, and cDNA31) (9). Compared to SARS-CoV-2 S protein, GD pCoV S proteins display several substitutions among different strains, including D290E, V320A, H519N/K, R634S, and S659T (**Fig. 1D**). Two substitutions (V320A and H519N/K) are in the RBD (25), while one substitution (D290E) is in the N-terminal domain of S (25). While some GD pCoVs (e.g. GD/M5-9/2019 and GD/P44-9/2019) contain incomplete or ambiguous sequences, no other nonsynonymous mutations or indels were observed (**Fig. S1**).

We then constructed the plasmids expressing the S proteins of GD pCoVs as well as SARS-CoV-2, and prepared lentivirus-based pseudoviruses harboring these S proteins. It should be noted that the S amino acid sequences were identical for cDNA8, cDNA9, cDNA20, and cDNA31; for GD/P79-9/2019 and cDNA16; and for GD/1/2019 and cDNA18 (**Fig. 1D**). Therefore, cDNA8, GD/1/2019, and GD/P79-9/2019 were used as representatives of the strains with redundant S sequences. Consistent with previous reports (20,21), GD/1/2019 and MP789 were capable of infecting human ACE2-expressing cells. Here we showed that all GD pCoVs can use human ACE2 as an infection receptor (**Fig. 1E**). The pseudovirus infectivity of GD/1/2019 (adjusted *P* < 0.0001), cDNA8 (adjusted *P* < 0.0001), GD/M5-9/2019 (adjusted *P* < 0.0001), MP789 (adjusted *P* < 0.0001), and GD/P44-9/2019 (adjusted *P* = 0.027) were significantly higher than that of SARS-CoV-2. On the other hand, the infectivity of GD/P79-9/2019 (adjusted *P* = 0.040) was significantly lower than that of SARS-CoV-2 (**Fig. 1E**).

To compare the susceptibility of GD pCoVs to antiviral humoral immunity elicited by SARS-CoV-2 vaccination, we performed neutralization assays using sera from individuals who received three doses of the Pfizer-BioNTech COVID-19 vaccine. As outlined in previous reports (22,23), GD/1/2019 is susceptible to anti-SARS-CoV-2 humoral immunity induced by SARS-CoV-2 vaccination (**Fig. 1F**). Here we showed that all GD pCoVs are sensitive to SARS-CoV-2 vaccine-induced humoral immunity (**Fig. 1F**). Notably, the sensitivity of GD pCoVs to vaccine sera differs between strains. While the 50% neutralization titer (NT50) of MP789 was comparable to that of SARS-CoV-2, vaccine sera exhibited significantly higher NT50s against GD/1/2019 (5.0-fold), cDNA8 (4.5-fold), GD/P79-9/2019 (4.8-fold), GD/M5-9/2019 (8.5-fold), and GD/P44-9/2019 (2.5-fold) compared to those against SARS-CoV-2 (**Fig. 1F**). Collectively, our results suggest that all GD pCoVs exhibited increased or comparable susceptibility to SARS-CoV-2 vaccine sera.

### Residue 519 is responsible for the increased sensitivity of some GD pCoVs to SARS-CoV-2 vaccine sera

To identify the amino acid residue(s) that determine the higher sensitivity of some GD pCoVs to SARS-CoV-2 vaccine sera, we again compared the amino acid sequences of GD pCoV S proteins. The GD pCoVs with higher sensitivity to vaccine sera (>4-fold; GD/1/2019, cDNA8, GD/P79-9/2019, and GD/M5-9/2019) all harbor asparagine (N) at residue 519 of S protein, while those with relatively comparable NT50 values to SARS-CoV-2 (MP789 and GD/P44-9/2019) harbor lysine (K) at this position (**Fig. 2A**, **Fig. S1**, and **Fig. S2**). In contrast to all GD pCoVs, SARS-CoV-2 S harbors histidine (H) at residue 519. To investigate the impact of residue 519 on the infectivity and immune susceptibility of GD pCoVs, we generated swapping substitutions at residue 519—the N519K derivatives for GD/1/2019, cDNA8, GD/P79-9/2019, and GD/M5-9/2019; the K519N derivatives for MP789 and GD/P44-9/2019; and the H519N and H519K derivatives for SARS-CoV-2.

**Fig. 2.**
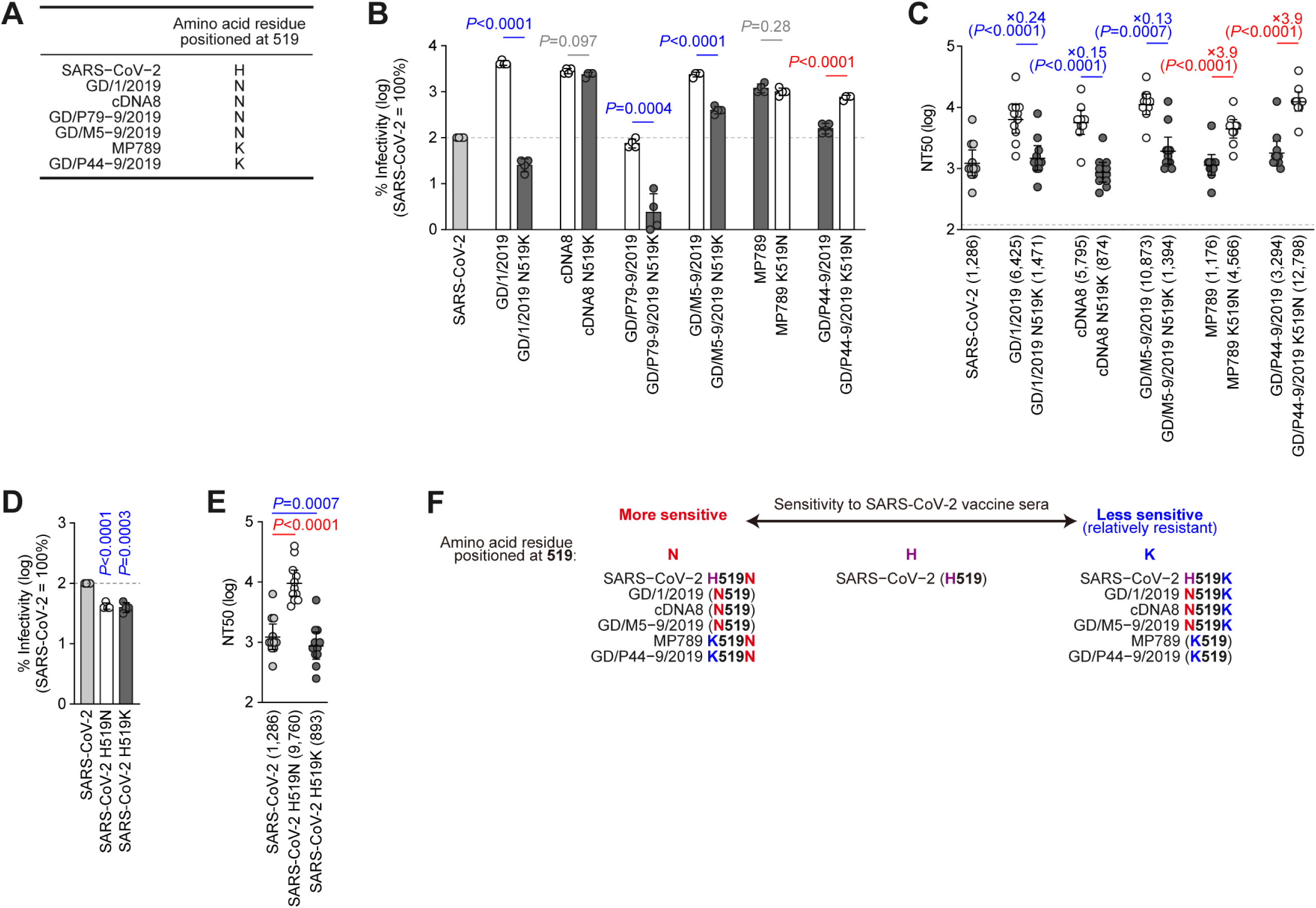
Effect of the amino acid substitution position at 519 on pseudovirus infectivity and sensitivity to SARS-CoV-2 vaccine sera. (**A**) Amino acid positioned at residue 519 in GD pCoVs and SARS-CoV-2. (**B**) Pseudovirus infection assay of GD pCoV 519 mutants. HIV-1-based reporter viruses harboring the S proteins of GD pCoVs and their derivatives were prepared. The pseudoviruses were inoculated into HOS-TMPRSS2 cells stably expressing human ACE2. (**C**) Neutralization assay of GD pCoV 519 mutants. Neutralization assays were performed with pseudoviruses harboring the S proteins of GD pCoVs, SARS-CoV-2, and their derivatives against sera elicited by 3-dose Pfizer-BioNTech COVID-19 vaccination (n = 11). Due to totally abrogated infectivity in the GD/P79-9/2019 N519K mutant, GD/P79-9/2019 was excluded in this neutralization assay. (**D**) Pseudovirus infection assay of SARS-CoV-2 519 mutants. HIV-1-based reporter viruses harboring the S proteins of SARS-CoV-2 and its derivative were prepared. The pseudoviruses were inoculated into HOS-TMPRSS2 cells stably expressing human ACE2. (**E**) Neutralization assay of SARS-CoV-2 519 mutants. Neutralization assays were performed with pseudoviruses harboring the S proteins of GD pCoVs, SARS-CoV-2, and their derivatives against sera elicited by 3-dose Pfizer-BioNTech COVID-19 vaccination (n = 11). (**F**) Schematic summarizing the effect that the amino acid positioned at residue 519 (i.e. asparagine; N, histidine; H, or lysine; K) has on the sensitivity of pseudoviruses to vaccine sera. In (**B**) and (**D**), data are expressed as the mean and SD of the log-transformed percent pseudovirus infectivity. The horizontal dashed line indicates the infectivity of SARS-CoV-2. Assays were performed in quadruplicate. Statistically significant differences versus parental strains (**B**) or SARS-CoV-2 (**D**) were determined by the two-sided Student’s t test and *P* values are shown in the figure. Red and blue values indicate increased and decreased by the 519 substitutions, respectively. In (**C**) and (**E**), assays for each serum sample were performed in triplicate to determine the 50% neutralization titer (NT50). Each point represents one NT50 value, and the geometric mean and 95% confidence intervals are shown. The number in parentheses indicates the geometric mean of NT50 values. The horizontal dashed line indicates the detection limit (120-fold). Statistically significant differences versus parental strains (**C**) or SARS-CoV-2 (**E**) were determined by the two-sided Student’s t-test paired by serum samples and *P* values are shown in the figure. Red and blue values indicate increased and decreased NT50s, respectively. The fold changes of NT50 versus parental are calculated as the average ratio of NT50s obtained from each pseudovirus. The fold changes versus parental are indicated with “X”. Information on the vaccine sera donors is summarized in **Table S3**. In (**B**)–(**E**), open circles/columns represent viruses that carry an asparagine at position 519, while filled circles/columns represent viruses that carry a lysine at position 519. SARS-CoV-2 is included as a control and is shaded in gray.

We then investigated how the residue 519 affects the infectivity of GD pCoVs. As shown in **Fig. 2B**, the infectivity of cDNA8 and MP789 was not significantly affected by the N/K swapping substitution at residue 519 in S. However, the N519K substitution conferred significantly decreased infectivity to GD/1/2019, GD/P79-9/2019, and GD/M5-9/2019, with completely abrogated infectivity in GD/P79-9/2019 (**Fig. 2B**). In contrast, in the case of GD/P44-9/2019, the K519N substitution significantly increased infectivity compared to parental S pseudoviruses (**Fig. 2B**). These results suggest that residue 519 possibly modulates the pseudovirus infectivity of some GD pCoVs.

Next, we assessed the influence of residue 519 on the sensitivity of GD pCoVs to SARS-CoV-2 vaccine sera. As shown in **Fig. 2C**, the N519K substitution in GD/1/2019, cDNA8 and GD/M5-9/2019 resulted in significantly lower NT50s exhibited by vaccine sera compared to parental S pseudoviruses. In sharp contrast, in the case of MP789 and GD/P44-9/2019, the K519N substitution resulted in significantly higher NT50s compared to parental S pseudoviruses (**Fig. 2C**). These results suggest that N519 confers higher sensitivity to vaccine sera, while K519 pseudoviruses exhibit lower immune sensitivity compared to pseudoviruses harboring parental S.

In the case of SARS-CoV-2 S, the amino acid substitutions at position 519 (i.e. H519N and H519K) both significantly decreased pseudovirus infectivity (**Fig. 2D**). However, vaccine sera displayed significantly higher (7.6-fold) NT50s against the H519N derivative compared to parental SARS-CoV-2 (**Fig. 2E**). On the other hand, the H519K derivative exhibited significantly lower NT50s compared to parental SARS-CoV-2 (**Fig. 2E**). Altogether, these results suggest that the sensitivity to SARS-CoV-2 vaccine sera of both GD pCoV and SARS-CoV-2 S pseudoviruses can be modulated by the residue 519 (i.e. N or K) of S protein (**Fig. 2F**).

### Residue 519 substitution modulates the sensitivity to monoclonal antibodies recognizing its epitope

To address the question of how residue 519 modulates immune evasion, we aligned the S structures of SARS-CoV-2, MP789, and GD/1/2019. We utilized MP789 and GD/1/2019 as representatives of GD pCoVs harboring lysine and asparagine respectively at position 519. Residue 519 is located at the base of the RBD (residues 319-541 of the S protein) and is away from the ACE2 receptor-binding motif (RBM; residues 438-506 of the S protein), a region in the RBD that forms a binding interface with the ACE2 receptor (25) (**Fig. 3A** and **3B**). Additionally, the position of residue 519 in the context of the RBD structure is comparable between SARS-CoV-2 (H519), MP789 (K519), and GD/1/2019 (N519) (**Fig. 3B**).

**Fig. 3.**
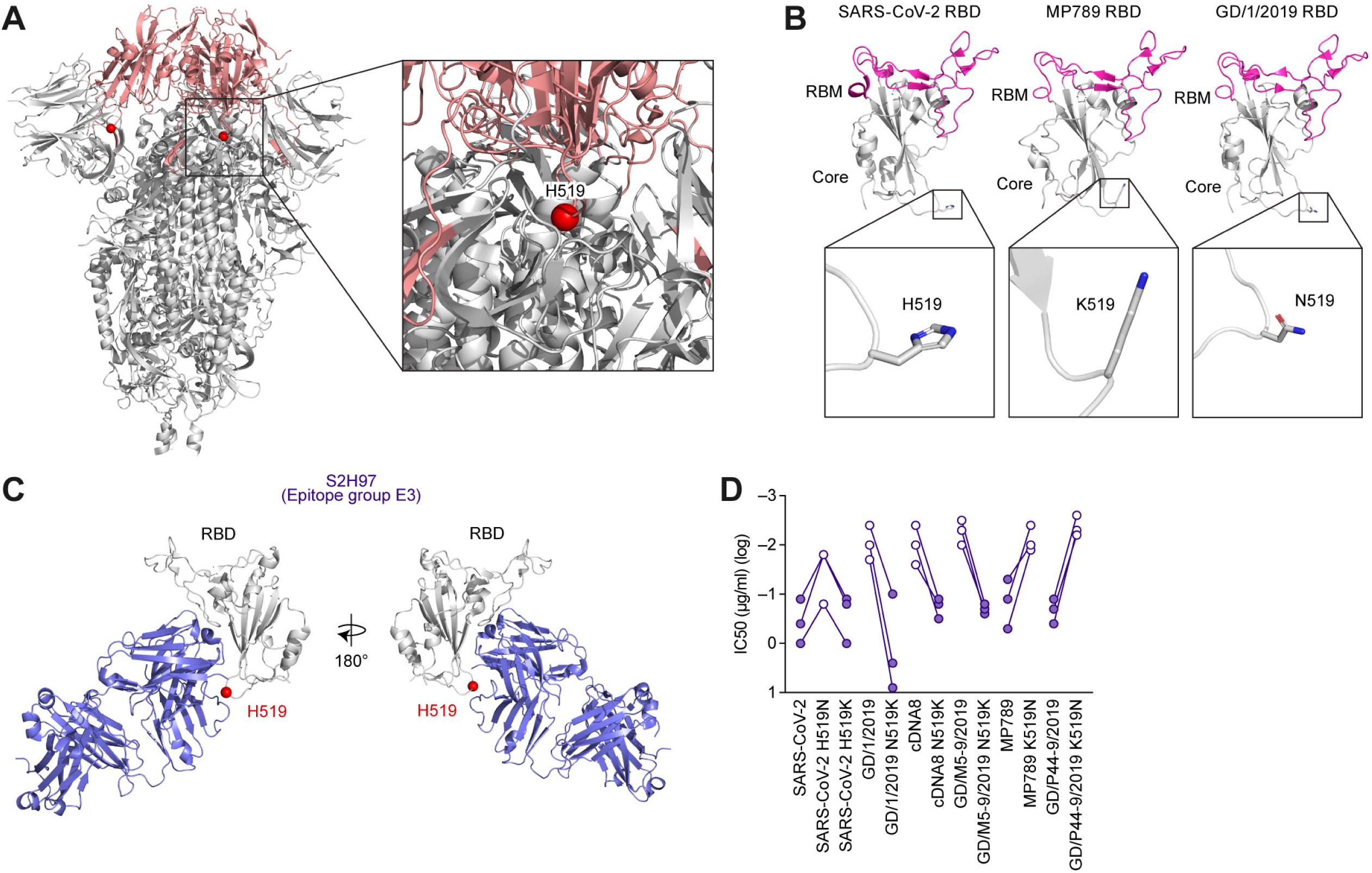
Effect of the amino acid substitution position at 519 on evasion from monoclonal antibodies targeting the epitope region including residue 519. (**A**) (left) The cryo-EM structure of the SARS-CoV-2 S trimer. The complex structure is shown as a cartoon. (right) The close-up view of the complex structure. The S RBD is highlighted in red and residue 519 is highlighted as a red sphere (PDB:6VXX). H519, histidine 519. (**B**) (top) The crystal structure of SARS-CoV-2 S RBD (PDB:6M0J), cryo-EM structure of MP789 S RBD (PDB:7BBH), and cryo-EM structure of GD/1/2019 RBD (PDB:7CN8). (bottom). Close-up views of the complex structures. Complex structures are shown as cartoons, and the residue positioned at 519 is shown as sticks. RBM, receptor-binding motif. (**C**) (left) Crystal structure of SARS-CoV-2 S RBD and S2H97 (PDB:7M7W). The complex structure is shown as a cartoon, and residue 519 is shown as a red sphere. (right) The structure rotated 180° on the y axis. RBD, receptor-binding domain. (**D**) Neutralization assay using three monoclonal antibodies recognizing epitope group E3. Neutralization assays were performed with pseudoviruses harboring the S proteins of GD pCoVs, SARS-CoV-2, and their derivatives against three epitope group E3-targeting mAbs (BD55-3337, BD55-5415, BD55-5583). Due to totally abrogated infectivity in the GD/P79-9/2019 N519K mutant, GD/P79-9/2019 was excluded in this neutralization assay. Assays for each antibody were performed in triplicate to determine the 50% inhibitory concentration (IC50). Each point represents one log-transformed IC50 value, and IC50s of the same antibodies are connected by a solid line for derivatives of the same virus. Open circles represent viruses that carry an asparagine at position 519, while filled circles represent viruses that carry a lysine or histidine at position 519. Exact IC50 values are summarized in **Table 1**.

**Table 1.** GD pCoV and SARS-CoV-2 neutralization by monoclonal antibodies.

| Antibody ID | Epitope Group <sup>a</sup> | ACE2 Competition <sup>a</sup> | IC50 (µg/ml) |  |  |  |  |  |  |  |  |  | SARS-CoV-2 | SARS-CoV-2 H519N | SARS-CoV-2 H519K |
| --- | --- | --- | --- | --- | --- | --- | --- | --- | --- | --- | --- | --- | --- | --- | --- |
|  |  |  | GD/1/2019 | GD/1/2019 N519K | cDNA8 | cDNA8 N519K | MP789 | MP789 K519N | GD/M5-9/2019 | GD/M5-9/2019 N519K | GD/P44-9/2019 | GD/P44-9/2019 K519N |  |  |  |
| BD55-3337 | E3 | 0.03 | 0.010 | 2.457 | 0.010 | 0.308 | 0.504 | 0.012 | 0.005 | 0.224 | 0.362 | 0.005 | 1.016 | 0.150 | 1.073 |
| BD55-5415 | E3 | 0.01 | 0.020 | 8.726 | 0.028 | 0.172 | 0.048 | 0.011 | 0.009 | 0.212 | 0.218 | 0.007 | 0.115 | 0.017 | 0.122 |
| BD55-5583 | E3 | 0.01 | 0.004 | 0.097 | 0.004 | 0.123 | 0.133 | 0.004 | 0.003 | 0.159 | 0.113 | 0.002 | 0.444 | 0.017 | 0.146 |
| BD30-503 | A | 0.86 | 0.006 | 0.011 | 0.010 | 0.017 | 0.031 | 0.013 | 0.008 | 0.009 | 0.017 | 0.005 | 0.015 | 0.015 | 0.028 |
| BD30-504 | A | 0.62 | 0.022 | 0.016 | 0.009 | 0.053 | 0.041 | 0.010 | 0.034 | 0.036 | 0.033 | 0.020 | 0.032 | 0.024 | 0.056 |
| BD55-300 | A | 0.92 | 0.004 | 0.026 | 0.004 | 0.005 | 0.011 | 0.004 | 0.004 | 0.019 | 0.009 | 0.004 | 0.008 | 0.006 | 0.008 |
| BD30-605 | A | 0.57 | 0.019 | 0.045 | 0.020 | 0.033 | 0.043 | 0.038 | 0.048 | 0.041 | 0.045 | 0.044 | 0.063 | 0.045 | 0.051 |
| BD55-5463 | B | 0.74 | 0.005 | 0.007 | 0.006 | 0.013 | 0.014 | 0.006 | 0.006 | 0.007 | 0.007 | 0.005 | 0.131 | 0.051 | 0.077 |
| BD55-203 | B | 0.8 | 0.014 | 0.023 | 0.011 | 0.041 | 0.050 | 0.011 | 0.018 | 0.047 | 0.053 | 0.010 | 0.038 | 0.011 | 0.011 |
| BD55-5966 | B | 0.89 | 0.002 | 0.033 | 0.003 | 0.004 | 0.006 | 0.005 | 0.003 | 0.007 | 0.006 | 0.003 | 0.019 | 0.005 | 0.009 |
| BD55-6382 | B | 0.82 | 0.004 | 0.040 | 0.003 | 0.004 | 0.011 | 0.004 | 0.003 | 0.007 | 0.006 | 0.003 | 0.011 | 0.003 | 0.005 |
| BD55-1239 | F2 | 0.82 | 0.022 | 0.018 | 0.016 | 0.034 | 0.122 | 0.020 | 0.028 | 0.119 | 0.109 | 0.027 | 0.217 | 0.037 | 0.181 |
| BD55-3500 | F2 | 0.91 | 0.022 | 0.034 | 0.028 | 0.034 | 0.064 | 0.034 | 0.024 | 0.066 | 0.067 | 0.014 | 0.343 | 0.064 | 0.393 |
| BD55-5380 | F2 | 0.81 | 0.009 | 0.044 | 0.006 | 0.009 | 0.031 | 0.016 | 0.022 | 0.026 | 0.022 | 0.015 | 0.154 | 0.004 | 0.093 |
| BD55-5189 | F2 | 0.77 | 0.008 | 0.047 | 0.028 | 0.130 | 0.076 | 0.013 | 0.020 | 0.082 | 0.069 | 0.016 | 0.069 | 0.007 | 0.035 |
| BD55-4637 | F3 | 0.81 | 0.019 | 0.019 | 0.013 | 0.077 | 0.062 | 0.013 | 0.015 | 0.077 | 0.039 | 0.015 | 0.103 | 0.031 | 0.090 |
| BD55-3372 | F3 | 0.88 | 0.004 | 0.020 | 0.004 | 0.009 | 0.012 | 0.005 | 0.009 | 0.018 | 0.018 | 0.009 | 0.006 | 0.006 | 0.012 |
| BD30-449 | F3 | 0.75 | 0.109 | 0.470 | 0.108 | 0.410 | 0.426 | 0.144 | 0.013 | 0.034 | 0.025 | 0.013 | 0.197 | 0.008 | 0.183 |
| BD55-5472 | F3 | 0.73 | 0.004 | 0.040 | 0.015 | 0.021 | 0.053 | 0.011 | 0.010 | 0.026 | 0.019 | 0.010 | 0.125 | 0.004 | 0.103 |

<sup>a</sup> Epitope Group and ACE2 competition levels were determined by Cao et al. (26). Their findings for the mAbs utilized in this study are reported here.

Next, we investigated the process by which residue 519 substitution impacts neutralization in SARS-CoV-2 and GD pCoVs. Since polyclonal vaccinated sera contain a mixture of multiple neutralizing antibodies, we utilized individual monoclonal antibodies (mAbs) against S RBD to characterize the effect of 519 substitution on different classes of neutralizing antibodies. In a deep mutational scanning screen of 1,538 mAbs against SARS-CoV-2 S pseudoviruses, Cao et al. (26) described 12 clusters of mAbs based on their escape mutations and dubbed them “epitope groups”. According to this definition, the residue 519 is in the epitope group E3. To evaluate the effect of residue 519 substitution on the immunological phenotype of epitope group E3, we used three mAbs targeting this epitope. As shown in **Fig. 3C**, S2H97 (PDB:7M7W) (27), which is a classical mAb targeting epitope group E3, recognizes the structural region including residue 519 (26). As expected, the neutralization assay using these three mAbs showed that the pseudoviruses harboring N519 exhibited lower 50% inhibitory concentrations (IC50s) than those harboring K519 (**Fig. 3D**). Altogether, these results suggest that N519 confers higher sensitivity to the mAbs targeting the epitope group E3, while K519 pseudoviruses exhibit lower sensitivity to these mAbs.

### Residue 519 substitution indirectly affects neutralization activity by up-state RBD-binding monoclonal antibodies

We showed that the neutralization activity of antibodies recognizing epitope group E3 was dramatically improved by asparagine substitution at residue 519 (**Fig. 3D**). However, the neutralization activity of these mAbs has been reported to be relatively weak across a broad spectrum of SARS-CoV-2 variants (26). Moreover, since vaccinated sera contain a variety of neutralizing antibodies, E3-targeting mAbs alone may not fully explain the magnitude of immune sensitivity displayed by N519-harboring pseudoviruses. Therefore, we assumed that the substitution of residue 519 might also affect the neutralization activity of other mAbs in a different fashion.

The RBD of both GD pCoVs and SARS-CoV-2 can conformationally shift into an open (“up”) conformational state, with potent antibodies binding to sites that become more exposed in the up-state RBD conformation (28,29). In fact, a previous study has suggested that residue 519 possibly modulates the up/down-state of the RBD (30). We therefore hypothesized that N519 may stabilize the up-state RBD conformation, consequently resulting in higher immune sensitivity to potent mAbs binding to sites exposed in the up-state RBD conformation. To investigate this hypothesis, we used mAbs targeting epitope groups A, B, F2, and F3 as described by Cao et al. (26). These mAbs bind at or near the RBM and compete with ACE2 by binding the up-state RBD conformation (**Table 1**) (26). Although residue 519 is not at the binding interface of these antibodies (**Fig. 4A**), we revealed that most of the up-state RBD-targeting mAbs exhibited higher IC50 values against the N519K derivatives of GD/1/2019 (13 out of 16), cDNA8 (15 out of 16), and GD/M5-9/2019 (14 out of 16) compared to parental S pseudoviruses (**Fig. 4B** and **Table 1**). In contrast, these mAbs exhibited lower IC50 values against the K519N derivatives of MP789 (15 out of 16) and GD/P44-9/2019 (15 out of 16) compared to parental S pseudoviruses (**Fig. 4B** and **Table 1**). In the case of SARS-CoV-2, H519N led to all 16 of the up-state RBD-targeting mAbs exhibiting lower IC50 values (**Fig. 4C** and **Table 1**). Collectively, our results suggest that N519 indirectly affects the conformation of RBD, thereby enhancing neutralization by up-state RBD-binding neutralizing antibodies in GD pCoVs and SARS-CoV-2 (**Fig. 4D**).

**Fig. 4.**
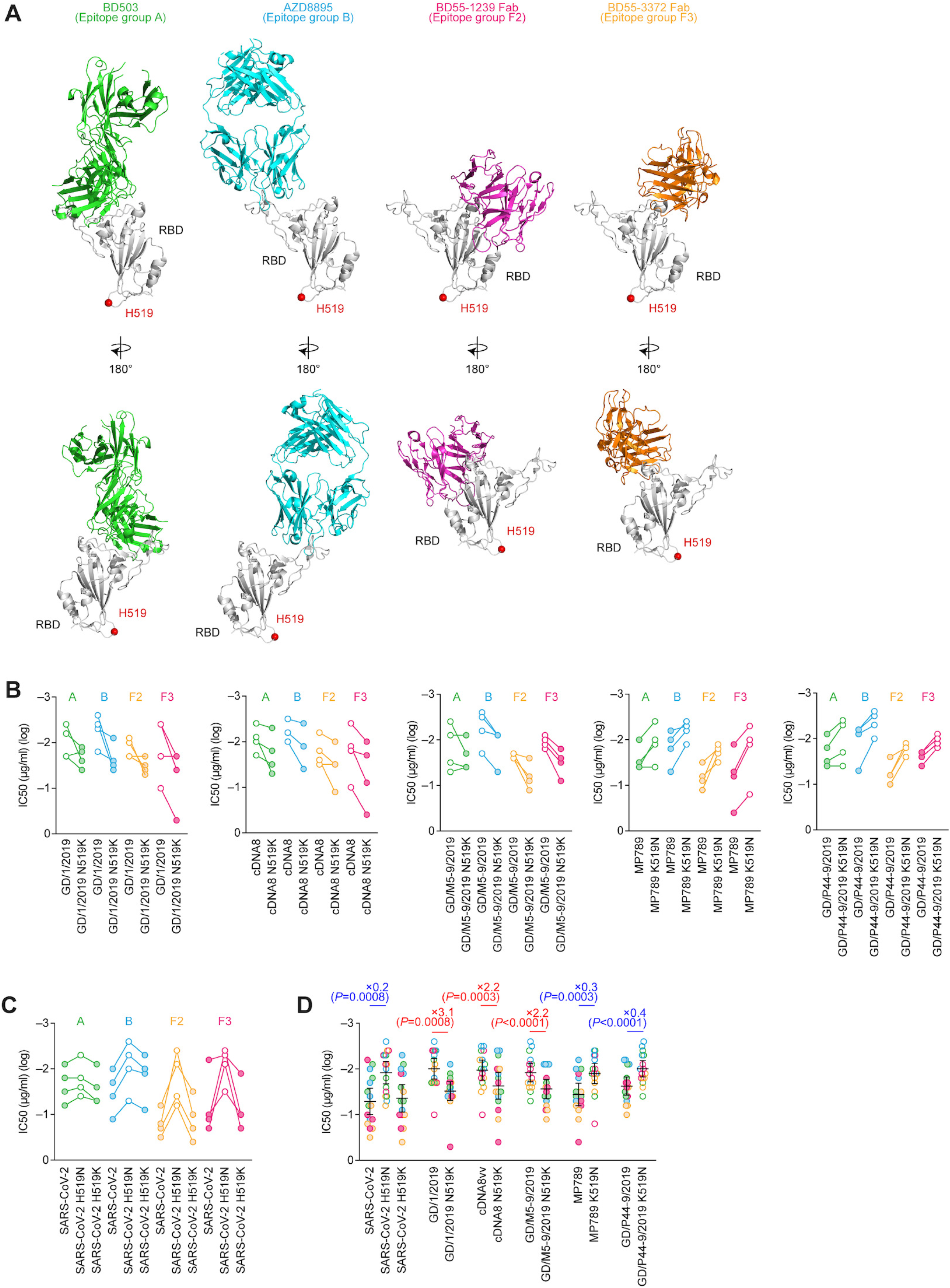
Effect of the amino acid substitution position at 519 on evasion from monoclonal antibodies potentially recognizing up-state RBD. (**A**) (top) Structural model of SARS-CoV-2 S RBD in complex with BD-503 (PDB:7EJY), AZD8895 (PDB:7L7D), BD55-1239 (PDB:7WRL), and BD55-3372 (PDB:7WRO). The complex structure is shown as a cartoon, and residue 519 is shown as a red sphere. (bottom) The structures rotated 180° on the y axis. RBD, receptor-binding domain. (**B–D**) Neutralization assay using monoclonal antibodies recognizing epitope groups A, B, F2 and F3. Neutralization assays were performed with pseudoviruses harboring the S proteins of GD pCoVs, SARS-CoV-2, and their derivatives against a total of 16 monoclonal antibodies recognizing epitope groups A, B, F2, F3. Assays for each antibody were performed in triplicate to determine the 50% inhibitory concentration (IC50). Due to totally abrogated infectivity in the GD/P79-9/2019 N519K mutant, GD/P79-9/2019 was excluded in this neutralization assay. Each point represents one log-transformed IC50 value exhibited by the antibodies against GD pCoVs (**B**) and SARS-CoV-2 (**C**). Plots are separated by parental strain, and in each plot, the points are color-coded by epitope group. IC50s of the same antibodies are connected by a solid line for derivatives of the same virus. Open circles represent viruses that carry an asparagine at position 519, while filled circles represent viruses that carry a lysine or histidine at position 519. Exact IC50 values are summarized in Table 1. Statistical comparisons of the data shown in (**B**) and (**C**) are summarized in (**D**). The geometric mean and 95% confidence intervals are shown. Statistically significant differences versus parental were determined by the two-sided Student’s t-test paired by antibody and *P* values are shown in the figure. Red and blue values indicate increased and decreased IC50s, respectively. The fold changes of IC50 versus parental are calculated as the average ratio of IC50 obtained from each pseudovirus. The fold changes versus parental are indicated with “X”. Exact IC50 values are summarized in **Table 1**.

## Discussion

Previous studies involving pCoVs have utilized only either GD/1/2019 (21,23,31) or MP789 (20) as a representative. In this study, we elucidate the previously unknown phenotypic diversity in all GD pCoVs known to date. We showed that the neutralization sensitivity of GD pCoVs differs between strains. Furthermore, we experimentally demonstrated that these differences are determined by the amino acid residue positioned at 519. Importantly, the impact of residue 519 is not restricted to GD pCoVs but is also observed in SARS-CoV-2.

The evolutionary implications of residue 519 substitution are striking in the context of the broader genetic landscape of sarbecoviruses. Asparagine at position 519 (N519) is remarkably conserved in most sarbecoviruses, and the S proteins of all bat sarbecoviruses analyzed in this study harbored N519 (**Fig. S2**). Interestingly, certain GD pCoVs harbor K519, and SARS-CoV-2 harbors H519. These observations suggest that SARS-CoV-2-related coronaviruses seem to prefer substituting residue 519 once out of their bat hosts. Because residue 519 possibly modulates the up/down-state of the RBD (30), conformational switching by residue 519 substitution may be critical for sarbecoviruses to spill over into non-bat hosts including pangolins and humans.

Since we find that K519 drives immune evasion, the evasion of antiviral humoral immunity in pangolins may have been a key factor driving the acquisition of K519 in some GD pCoVs. Supporting this idea, Wacharapluesadee et al. (15) have detected antibodies that cross-react with SARS-CoV-2 in pangolin serum in Southern Thailand. Since the serological and/or virological prevalence of pCoVs in wild pangolins in Southeast Asia remains unclear, it is unknown whether there is pre-existing anti-pCoV immunity in wild pangolins. However, it might be possible to assume that K519 substitution was acquired to evade pre-existing anti-pCoV humoral immunity in wild pangolins. Since early SARS-CoV-2 sequences all harbor H519 (32), this also raises the critical question of whether the substitution from asparagine to histidine occurred in a putative intermediate host or shortly after the emergence of SARS-CoV-2 in humans.

In addition to differences in immune susceptibility, pseudovirus infectivity was also different between the GD pCoV strains: the N519K substitution significantly decreased pseudovirus infectivity in some N519 viruses (GD/1/2019, GD/P79-9/2019, GD/M5-9/2019) and *vice versa* (K519N; GD/P44-9/2019) but did not significantly affect the pseudovirus infectivity of cDNA8 and MP789 (**Fig. 2B**). Therefore, the difference in pseudovirus infectivity among GD pCoV strains cannot be determined by residue 519 alone (**Fig. 1E**). The effect of residue 519 on viral infectivity may depend on epistatic interactions with other residues, including those located in the backbone S protein.

In conclusion, our study explored a hereto unknown phenotypic diversity within the pCoVs detected in the Guangdong province of China. We demonstrated that substitutions within GD pCoVs contribute to differences in the infectivity and immune sensitivity of these viruses. Notably, we identified a substitution from asparagine to lysine at position 519 as a key driver of lowered immune sensitivity. This suggests that lysine substitution at position 519 is a potent immune escape mutation harbored by GD pCoVs including MP789 and GD/P44-9/2019. Strikingly, MP789 not only has relatively lower immune sensitivity to vaccine sera (**Fig. 1F**) but also has greater infectivity than SARS-CoV-2 (**Fig. 1E**). Therefore, further surveillance of pCoVs, especially those harboring K519, should be a key component of measures to anticipate and mitigate future outbreaks. Nevertheless, it should be noted that SARS-CoV-2 vaccine sera are still capable of satisfactorily neutralizing all GD pCoVs. The intricacies of the immunity of bats, pangolins, putative intermediate hosts, and humans, which make it evolutionarily advantageous for residue 519 substitutions to occur in non-bat hosts, should also be further explored.

## Methods

### Ethics statement

All protocols involving specimens from human subjects recruited at Kyoto University were reviewed and approved by the Institutional Review Boards of Kyoto University (approval ID: G1309). All human subjects provided written informed consent. All protocols for the use of human specimens were reviewed and approved by the Institutional Review Boards of The Institute of Medical Science, The University of Tokyo (approval IDs: 2021-1-0416 and 2021-18-0617) and Kyoto University (approval ID: G0697).

### Nucleotide sequence data collection

The nucleotide sequences of sarbecoviruses with *Manis javanica* as their host were retrieved from NCBI virus (download date, November 11, 2024). In addition, SARS-CoV-2 (Wuhan-Hu-1) and relevant sarbecovirus sequences were collected from NCBI GenBank (download date, November 11, 2024). GD/1/2019 (9) was accessed from GISAID on February 6, 2025 (accession number EPI_ISL_410721). Information on the sarbecovirus sequences is summarized in **Table S1**.

### Phylogenetic analysis

To evaluate the phylogenetic relationships between all sarbecoviruses, we aligned the complete genomes using the *-localpair* option of MAFFT v7.526 (33). The aligned sequence was cleaned using an in-house python script to replace any characters not in ‘ATGCN-’ with ‘N’. Next, we inferred a maximum likelihood (ML) tree based on this alignment using IQ-TREE2 v2.2.6 (34) under a GTR+F+I+R4 substitution model (**Fig. 1A**). Node support was assessed with 1000 Ultrafast bootstrap replicates (35). Next, we extracted the S-encoding part of the genomes (using the SARS-CoV-2 Wuhan-Hu-1 strain S protein as a reference) and used the method above to construct an S tree based on these nucleotide sequences (**Fig. 1B**). Based on the S sequences, we manually extracted the RBD sequences using the SARS-CoV-2 RBD as a reference (i.e. amino positions 319-541 in the SARS-CoV-2 Wuhan-Hu-1 strain S protein) and constructed an amino acid alignment using MAFFT v7.526 (33). This alignment was used to create an ML tree using IQ-TREE2 v2.2.6 (34) under the best substitution model selected by ModelFinder (36) with 1000 bootstrap iterations.

### Amino acid sequence alignment

To assess the similarity between all GD pCoVs, we utilized the sequences of SARS-CoV-2 Wuhan-Hu-1, GD/1/2019, cDNA8, cDNA9, cDNA16, cDNA18, cDNA20, cDNA31, MP789, GD/P79-9/2019, GD/M5-9/2019, and GD/P44-9/2019. Sequence information is summarized in **Table S1**. We extracted the S-encoding part of the genomes (using the SARS-CoV-2 Wuhan-Hu-1 strain S protein as a reference) for the GD pCoV sequences above and constructed an amino acid sequence alignment using MAFFT v7.526 (33) with the GTR+F+I+R4 substitution model, using 1000 maximum iterative refinements. Then, the substitutions within the GD pCoV sequences were manually confirmed in reference to the SARS-CoV-2 Wuhan-Hu-1 amino acid sequence (**Fig. S1**).

### Plasmid construction

Plasmids expressing the codon-optimized S proteins of GD pCoVs were synthesized by a gene synthesis service (Fasmac). The plasmid expressing the codon-optimized SARS-CoV-2 S protein (strain Wuhan-Hu-1; GenBank accession number: NC_045512.2) (32) was kindly provided by Dr. Kenzo Tokunaga. Plasmids expressing the residue 519 swapping substitutions of the S proteins of SARS-CoV-2 and GD pCoVs were generated by site-directed overlap extension PCR using the templates and primers listed in **Table S2**. The resulting mutagenized PCR fragments were cloned into the KpnI/NotI site of the pCAGGS vector (37) using the In-Fusion HD Cloning Kit (Takara, Cat# Z9650N). Nucleotide sequences were determined by DNA sequencing services (Eurofins), and the sequence data were analyzed by SnapGene v8.0.1 (SnapGene software).

### Cell culture

Lenti-X 293T cells (a human embryonic kidney cell line; ATCC, Takara, Cat# 632180) and HOS-ACE2/TMPRSS2 cells (kindly provided by Dr. Kenzo Tokunaga), a derivative of HOS cells (a human osteosarcoma cell line; ATCC CRL-1543) stably expressing human ACE2 and TMPRSS2 (38,39) were maintained in Dulbecco’s modified Eagle’s medium (high glucose) (Sigma-Aldrich, Cat# 6429-500ML) containing 10% fetal bovine serum (Sigma-Aldrich Cat# 172012-500ML) and 1% penicillin-streptomycin (Sigma-Aldrich, Cat# P4333-100ML).

### Pseudovirus assay

Pseudoviruses were prepared as previously described (40–44). Briefly, lentivirus (HIV-1)-based luciferase-expressing reporter viruses were pseudotyped with the S proteins of GD pCoVs or SARS-CoV-2 and their derivatives. LentiX-293T cells (500,000 cells) were cotransfected with 0.8 μg psPAX2-IN/HiBiT (38), 0.8 μg pWPI-Luc2 (45), and 0.4 μg plasmids expressing parental S or its derivatives using TransIT-293 (Takara, Cat# MIR2704) according to the manufacturer’s protocol. Two days post transfection, the culture supernatants were harvested, and the pseudoviruses were stored at –80°C until use. For pseudovirus infection, the amount of input virus was normalized to the HiBiT value measured by the NanoGlo HiBiT lytic detection system (Promega, Cat# N3040) as previously described (45). In this system, HiBiT peptide is produced with HIV-1 integrase and forms NanoLuc luciferase with LgBiT, which is supplemented with substrates. In each pseudovirus particle, the detected HiBiT value is correlated with the amount of the pseudovirus capsid protein, HIV-1 p24 protein (45). Based on the HiBiT value measured, we calculated the amount of HIV-1 p24 capsid protein according to the previous paper (45). In the pseudovirus infection assays, pseudoviruses were normalized to 4 ng of HIV-1 p24 capsid protein. At two days post infection, the infected cells were lysed with a BrightGlo luciferase assay system (Promega, Cat# E2620), and the luminescent signal produced by firefly luciferase reaction was measured using a GloMax Explorer Multimode Microplate Reader (Promega).

### Human serum collection

Vaccine sera were collected from individuals who had received three doses of the Pfizer-BioNTech COVID-19 vaccine (time interval between the last vaccination and sampling: 16-33 days; n=11; average age: 39.0 years, range: 29-53 years, 36.4% male). Sera were inactivated at 56°C for 30 minutes and stored at –80°C until use. The details of the vaccine sera are summarized in **Table S3**.

### Antibody isolation and recombinant production

The 19 monoclonal antibodies used in this study (listed in **Table 1**) were kindly provided by Dr. Yunlong Cao. These antibodies were generated as previously described by Cao et al. (26).

### Neutralization assay

Pseudovirus neutralization assays against vaccine sera and mAbs were performed as previously described (40–42,46). Briefly, the GD pCoV and SARS-CoV-2 S pseudoviruses (counting ∼100,000 relative light units) were incubated with serially diluted (120-fold to 87,480-fold dilution at the final concentration) heat-inactivated sera or serially diluted (0.008-fold to 125-fold the IC50 reported by Cao et al. (26) at the final concentration) mAbs at 37°C for 1 hour. Pseudoviruses without sera or mAbs were included as controls. Then, a 20 µl mixture of pseudovirus and serum/antibody was inoculated into HOS-TMPRSS2 cells stably expressing human ACE2 (10,000 cells/100 µl) in a 96-well white plate. Two days post infection, the infected cells were lysed with a Bright-Glo luciferase assay system (Promega, Cat# E2620) and the luminescent signal was measured using a GloMax explorer multimode microplate reader 3500 (Promega). The assay of each serum sample was performed in triplicate, and the 50% neutralization titer (NT50) or the 50% inhibitory concentration (IC50) was calculated using Prism 10 software v10.3.1 (GraphPad Software).

### Protein structure model

In **Fig. 3A**, the Cryo-EM structure of the S trimer of SARS-CoV-2 was used (PDB:6VXX) (47). In **Fig. 3B**, the crystal co-structure of SARS-CoV-2 S (PDB:6M0J) (25), cryo-EM structure of MP789 S (PDB:7BBH) (20), and cryo-EM structure of GD/1/2019 S (PDB:7DDO) (21) were used. In **Fig. 3C**, the structure of S2H97 (PDB:7M7W) (27) was used. In **Fig. 4A**, the structure of BD-503 (PDB:7EJY) (48), structure of AZD8895 (PDB: 7L7D) (49), structure of BD55-1239 Fab (PDB:7WRL) (26), and the structure of BD55-3372 Fab (PDB:7WRO) (27) were used. All protein structural analyses were performed using the PyMOL molecular graphics system v3.0.0 (Schrödinger).

## Statistical analysis

Statistical significance in the pseudovirus infectivity and neutralization assay was tested using the two-sided Student’s t-test on log-transformed data. All *P* values were adjusted to account for multiple comparisons using the Holm-Šídák method. For the neutralization assays, NT50 and IC50 data from the same serum samples and mAbs respectively were paired. The tests above were performed using Prism 10 software v10.3.1 (GraphPad Software). *P* < 0.05 was considered significant.

## Data and code availability

All databases/datasets used in this study are available from the GenBank database (https://www.ncbi.nlm.nih.gov/genbank/) and the GISAID database (https://www.gisaid.org; EPI_ISL_410721). All raw files used in this study are available on the GitHub repository (https://github.com/TheSatoLab/pCoV_519). Any additional information required to reanalyze the data reported in this work is available from the lead contact upon request.

## Figure legend

**Fig. S1 Amino acid sequence alignment of SARS-CoV-2 and GD pCoVs.** The amino acid residues between residues 285 to 300, 315 to 325, 515 to 525, 630 to 640, and 655 to 665 in SARS-CoV-2 and GD pCoVs. The residues different among pCoVs positioned at 290, 320, 519 and 634 are indicated.

**Fig. S2 Amino acid residue positioned at 519 in Sarbecoviruses.** Maximum likelihood trees of SARS-CoV-2 related pangolin coronaviruses, SARS-CoV-2 (strain Wuhan-Hu-1), and other sarbecoviruses based on the S RBD amino acid sequences. SARS-CoV-1 (strain Tor2) and two SARS-CoV-related coronaviruses (strains WIV1 and LYRa11) are included as an outgroup. Node support is shown above key branches. Scale bar indicates genetic distance (amino acid substitutions per site). The amino acid sequence alignment of the S protein from position 515 to position 525 is aligned to the tree. Sequences that do not harbor an asparagine at position 519 are highlighted in yellow. GD pCoVs and GX pCoVs are annotated based on previous published work (9–12).

**Table S1 Sequence data and accession numbers used in this study**.

**Table S2 Primers used in this study**.

**Table S3 Human 3-dose Pfizer-BioNTech vaccine sera used in this study.**

## Acknowledgements

We gratefully acknowledge all data contributors, i.e. the Authors and their Originating laboratories responsible for obtaining the specimens, and their Submitting laboratories for generating the genetic sequence and metadata and sharing via the GISAID Initiative, on which this research is based.

We would like to express our gratitude to all the members of Division of Systems Virology, The Institute of Medical Science, The University of Tokyo. We would like to express special thanks to Dr. Spyros Lytras for substantial and insightful comments on the manuscript.

This study was supported in part by AMED ASPIRE Program (24jf0126002, to G2P-Japan Consortium and Kei Sato); AMED SCARDA Japan Initiative for World-leading Vaccine Research and Development Centers “UTOPIA” (243fa627001, to Kei Sato); AMED SCARDA Program on R&D of new generation vaccine including new modality application (243fa727002, to Kei Sato); AMED Research Program on Emerging and Re-emerging Infectious Diseases (23fk0108583, JP24fk0108690, to Kei Sato); AMED Japan Program for Infectious Diseases Research and Infrastructure (Collaborative Research via Overseas Research Centers) (24wm0225041, to Kei Sato); JSPS KAKENHI Fund for the Promotion of Joint International Research (International Leading Research) (JP23K20041, to G2P-Japan Consortium and Kei Sato); JSPS KAKENHI Grant-in-Aid for Scientific Research A (JP24H00607, to Kei Sato); Mitsubishi UFJ Financial Group, Inc. Vaccine Development Grant (to Kei Sato); and The Cooperative Research Program (Joint Usage/Research Center program) of Institute for Life and Medical Sciences, Kyoto University (to Kei Sato); JSPS Research Fellow DC1 (23KJ0710, to Yusuke Kosugi); JSPS KAKENHI Grant-in-Aid for Scientific Research A (JP24H00607, to Kei Sato); Mitsubishi UFJ Financial Group, Inc. Vaccine Development Grant (to Kei Sato); Maximilian Stanley Yo is supported by a grant from the Japanese Government Ministry of Education, Culture, Sports, Science and Technology Scholarship– Research Category (240042). Jarel Elgin Tolentino is supported by a grant from the Japanese Government Ministry of Education, Culture, Sports, Science and Technology Scholarship–Research Category (220235).

